# Three-dimensional nano-imaging reveals subtle changes in xylem structure in CAD-deficient sorghum

**DOI:** 10.64898/2026.02.21.707186

**Authors:** Lais B. Manoel, Francine F. Fernandes, Eduardo Monteiro, Leydson G. A. Lima, Tiago A. Kalile, Florian Meneau, Igor Cesarino, Carla C. Polo

## Abstract

Lignin plays a central role in the formation and function of secondary cell walls in vascular plants. However, the structural consequences of lignin modification for cell wall properties and cellular function in grasses remain poorly understood. Here, we investigated how cinnamyl alcohol dehydrogenase (CAD) deficiency alters vascular cell architecture in *Sorghum bicolor*, using the brown midrib-6 (*bmr6*) mutant as a model system. Biochemical and histochemical analyses confirmed altered lignin chemistry in *bmr6*, including increased incorporation of hydroxycinnamaldehyde residues and reduced tricin levels. We applied ptychographic X-ray computed tomography (PXCT) to quantify the cell wall geometry, in three dimensions, at nanometer-scale resolution. PXCT enabled measurements of wall thickness distribution and lumen shape along tracheary elements. Analyses revealed no significant differences in wall thickness between wild-type and *bmr6* plants. However, three-dimensional morphometric descriptors indicated reduced lumen convexity in *bmr6*, suggesting localized modifications not detectable by conventional two-dimensional imaging. Water flow numerical simulations through PXCT-derived images indicated reduced vessel permeability and simulated hydraulic conductivity in *bmr6*, suggesting that subtle geometric changes may influence performance. These findings highlight the value of three-dimensional imaging for resolving cell wall organization and provide new insight into the architectural resilience of grass xylem in response to targeted lignin modification.

**Highlight:** Three-dimensional X-ray nano-imaging reveals alterations in the cell wall architecture that affect simulated hydraulic performance under reduced CAD activity in sorghum.

## Introduction

The plant cell wall is a dynamic structure that provides mechanical strength and structural support through its highly ordered architecture, thereby influencing cell morphology (Hoffmann *et al*., 2021). The cell wall is sufficiently flexible to resist shear forces but remains permeable enough to allow molecular transport (Quiroz-Castañeda and Folch-Mallol, 2011). Although the general architecture of the cell wall is conserved across angiosperms, its complexity is reflected by the large number of genes encoding proteins involved in its biosynthesis and assembly (Somerville et al., 2004). During the elongation phase, all plant cells are surrounded by a flexible primary wall composed of the structural polysaccharides cellulose, various hemicelluloses and pectins. Once elongation ceases, a thick secondary wall is deposited in specific cell types to allow functional specialization. For instance, secondary cell walls (SCW) are typically found in fibers, to bestow plants with physical support, and in tracheary elements of the xylem, to enable transport of water and nutrients throughout the plant body (Meents *et al*., 2018). SCWs consist of cellulose microfibrils, various hemicelluloses and the aromatic polymer lignin, and these components are present at varying proportions and different chemistries according to the cell type, taxon, and to responses to internal and environmental signals (Hoffmann *et al*., 2021). The differential impregnation of polysaccharides with lignin controls the hygroscopic capacity of the cell wall, fine-tuning the biochemical and biomechanical properties of cells and tissues to support their physiological functions (Pesquet *et al*., 2025).

Lignin is an insoluble, amorphous, and hydrophobic phenolic heteropolymer derived from the oxidative polymerization of different phenolic monomers, where the most abundant are the hydroxycinnamyl alcohols *p*-coumaryl, coniferyl, and sinapyl alcohol. The incorporation of these monolignols into the polymer occurs after oxidation by laccases and/or peroxidases, forming H-units (*p*-hydroxyphenyl), G-units (guaiacyl), and S-units (syringyl), respectively (Donaldson, 2001; Cesarino, 2019; Terrett and Dupree, 2019). In addition to the canonical monolignols, metabolites from different classes of the phenolic metabolism are also naturally incorporated into lignins of different tissues or plant species (Vanholme *et al*., 2019; Pesquet *et al*., 2025). For instance, the seed coat of some Cactaceae and Orchidaceae synthesizes lignins made exclusively of caffeyl alcohol (Chen *et al*., 2012, 2013), whereas lignin in grasses incorporates the flavone tricin (Lan *et al*., 2016). However, although different cell types have been described to synthesize chemical/structurally distinct lignin polymers, still little is known about how these different lignin chemistries affect the molecular, biochemical, and biomechanical properties of the cell wall and, consequently, how they contribute to the unique physiological roles of different specialized cell types (Pesquet *et al*., 2025).

The incorporation of alternative lignin monomers may contribute to elucidate the role of different lignin chemistries in cell wall properties and cell function. Among different strategies, these modifications can be achieved by misregulation of genes encoding lignin biosynthetic enzymes. In the lignin pathway, cinnamyl alcohol dehydrogenase (CAD) catalyzes the reduction of hydroxycinnamaldehydes into their corresponding hydroxycinnamyl alcohols (Sibout *et al*., 2005; Bouvier D’Yvoire *et al*., 2013; Ferreira *et al*., 2019), which is the last step in the biosynthesis of monolignols. CAD deficiency has often no impact on lignin content, whereas typically results in lignins enriched in coniferaldehyde and sinapaldehyde units, incorporated as cross-coupling moieties and end groups via characteristic aldehyde-derived β-aryl ether linkages (Zhao *et al*., 2013; Anderson *et al*., 2015; Ferreira *et al*., 2022). These structural modifications are known to improve sugar release from plant biomass upon industrial processing, although the exact mechanism leading to increased cell wall digestibility remains poorly understood. One hypothesis is that the strong dipoles contributed by the aldehyde groups may disrupt the intermolecular forces that facilitate integration of lignin with other cell wall biopolymers, increasing the accessibility of the carbohydrate components by hydrolytic enzymes (Anderson *et al*., 2015). Indeed, molecular dynamics simulation analysis found reduced non-covalent association between hemicellulose and aldehyde-rich lignins when compared to wild-type lignins (Vanholme *et al*., 2012). Additionally, as aldehyde units exhibit a higher propensity to terminate the lignin chains, the resulting polymer often shows lower molecular weights (Baumberger *et al*., 2002; Jourdes *et al*., 2007; Madigal *et al*., 2025) that, in turn, is expected to fail in protecting the structural polysaccharides as performed by larger polymers (Lapierre *et al*., 2004; Anderson *et al*., 2015). The observation that the level of aldehydes incorporation varies naturally among lignins from different cell types (Hänninen *et al*., 2011; Blaschek *et al*., 2020; Ménard *et al*., 2022) suggest an important biological function. Recent biomechanical experiments using different lignin mutants or transgenic lines in *Arabidopsis thaliana* and poplar showed that, for similar aromatic substitutions, lignin units with alcohol aliphatic functions increased stiffness whereas those with aldehyde functions increased flexibility of the cell wall (Özparpucu et al. 2017; Ménard et al. 2022). However, how the incorporation of aldehydes into lignin affects cell wall assembly and 3D architecture at the nanoscale remains largely uncharacterized.

Sorghum [*Sorghum bicolor* (L.) Moench] is a versatile annual crop cultivated as a source of grain, forage biomass, or sugar-rich stem juice but it is also considered a genetic model for bioenergy C4 grasses within the Panicoideae subfamily (Vermerris, 2011; van der Weijde *et al*., 2013). In addition to its C4 photosynthetic mechanism that promotes high biomass production, another advantage of sorghum for studies of biomass composition and digestibility is the availability of *brown midrib* (*bmr*) mutants, which show a characteristic brown coloration of the leaf midveins associated with reduced lignin content and/or altered lignin composition (Sattler *et al*., 2010). The *bmr6* mutant was originally isolated from diethyl sulfate-mutagenized populations and its causative gene was subsequently identified to encode SbCAD2 (Sobic.004G071000), the predominant CAD in sorghum (Saballos *et al*., 2009; Sattler *et al*., 2009). Further analyses of the *bmr6* mutant revealed reduced expression of *SbCAD2*, undetectable levels of its corresponding protein, and reduced CAD activity when compared with control plants (Sattler *et al*., 2009; Scully *et al*., 2016*a*). The *bmr6* mutant has been extensively characterized in terms of biomass composition and digestibility, showing decreased lignin levels (Saballos *et al*., 2008), lignin polymers enriched with hydroxycinnamaldehydes (Bucholtz *et al*., 1980; Palmer *et al*., 2008; Ferreira *et al*., 2022), altered S/G ratio (Sattler *et al*., 2009; Ferreira *et al*., 2022), and enhanced biomass digestibility (Godin *et al*., 2016; Scully *et al*., 2016 *a*). However, much less is known regarding the effects of reduced CAD activity (and the consequent alteration in lignin structure) on the cell wall architecture and functionality.

Synchrotron X-ray based microscopies (SXM), due to the short wavelength of X-rays and source brilliance, allows for accessing and analyzing the tissue volume to acquire morphological information of cell walls within their histological context, at high spatial resolution. Ptychographic X-ray computed tomography (PXCT) is a SXM technique developed at the 3^rd^ and 4^th^ synchrotron generations with full potential to explore three-dimensional organization of porous and hierarchical materials (Holler *et al*., 2017; Górecki *et al*., 2023; Báfero *et al*., 2026) and plant tissues/cells, such as tracheary elements, sclerenchymatic fibers, and parenchyma cells, revealing nano-scale morphological differences in cell walls that may arise from the misregulation of cell wall-related genes (Polo *et al*., 2020). Ultimately, this technique may contribute to not only our understanding of gene functions but also on how structural/chemical alterations in cell wall biopolymers affect the cell wall assembly, morphology and functionality. While optical microscopy is widely accessible and allows for the visualization of cellular structures in relatively large tissues, PXCT offers superior spatial resolution. PXCT enables volumetric measurements of cell wall thickness and lumen geometry along intact xylem elements at nanometer-scale resolution, providing structural information inaccessible to conventional two-dimensional imaging. Here, we investigate how lignin structural modification resulting from CAD deficiency affects the cell wall architecture of xylem cells in sorghum using high-resolution PXCT. We show that, despite substantial remodeling of lignin chemistry in *bmr6*, wall thickness of water conducting cells remains largely unchanged compared with wild-type plants, while subtle differences in three-dimensional lumen geometry and hydraulic conductivity are detectable. These findings demonstrate the structural resilience of the water conducting cell walls to targeted lignin modification and highlight the value of three-dimensional nano-imaging approaches for linking cell wall chemistry to architecture and cellular functions.

## Material and Methods

### Plant material and growth conditions

Seeds of sorghum wild-type (WT, CMSXS 101B) and *bmr6* mutant (CMSXS 101B *bmr6*) (Embrapa Milho e Sorgo - Sete Lagoas/MG) were germinated at the Brazilian Synchrotron Light Laboratory (LNLS) and grown in a greenhouse at the National Biorenewables Laboratory (LNBr), both located at the campus of the National Center for Research in Energy and Materials (CNPEM) in Campinas, São Paulo. The seedling process is summarized in Supplementary Figure S1. The seeds were sown in germination trays in a plant growth chamber (SGC 2®; Weiss Technik UK Ltd., Loughborough, UK), programmed for a photoperiod of 14 h:10 h light: dark, with temperature and humidity set at 25 °C and 60%, respectively. Thirty seeds from each genotype were planted in a substrate mixed with vermiculite (2:1). Irrigation was performed using an adapted system based on the pressure difference, where distilled water was released according to the plants’ evapotranspiration. After 30 days, the seedlings were transplanted into 14.3 L pots filled with substrate and vermiculite (2:1). Two seedlings were planted equidistantly per pot. In total, 12 WT individuals and 14 *bmr6* individuals were selected for cultivation in the greenhouse. According to the National Institute of Meteorology (INMET, 2024), during the period in the greenhouse, the average temperature was 23.1°C, the average radiation rate was 1480.4 (KJ/m²), and the relative humidity was 70.8% (estimates based on measurements from the Automatic Meteorological Station of Piracicaba, INMET, for the period from 17/01/2023 to 05/05/2023). Irrigation was carried out using an automatic drip system, set to remain activated for 10 min, four times a day, at 3 a.m., 9 a.m., 3 p.m., and 9 p.m. Harvesting of plant material was performed 110 days after transplanting, with a total growth period of 140 days.

### Morphological Characterization

The specimens were harvested 140 days post-cultivation, following the attainment of grain maturity by all panicles (May/2023). This timing was selected to ensure the full maturity of the plants for subsequent analyses. During the harvesting process, phenotypic characterization of each WT and *bmr6* plant was conducted. Prior to pruning, the substrate was partially removed to expose roots. The plant height was measured from the stem base to the panicle top. The total lengths of the panicles and flag leaves were recorded at this stage. The internodes were numbered in a base-to-apex direction, with internodes #3 and #4 selected for measurements. For biochemical and imaging analyses, internode samples were harvested. For each collected sample, the total length and diameter in the median region of internodes #3 and #4 were measured. Their central regions were isolated and sectioned into slices of approximately 1 cm³ for further analysis. During sectioning, priority was given to preserving the outer region near the rind, due to the presence of a higher number of lignified vascular bundles. The fragments were directly immersed in a fixative solution composed of formaldehyde, glacial acetic acid, and 70% ethyl alcohol (FAA70) in a volumetric ratio of 18:1:1. These samples were subjected to vacuum conditions for 24 h following the methodology outlined by Johansen (1940). Subsequently, the samples were preserved in 70% ethanol until histochemical analyses were conducted to histochemical assess lignin content. The morphological characterization considered 12 and 14 biological replicates of WT and *bmr6*, respectively and the differences between the genotypes were assessed by t-test with a significance threshold of *P <* 0.05.

### Lignin Analyses

Lyophilized internode material was subjected to a sequential extraction with water (98 °C), ethanol (76 °C), chloroform (59°C), and acetone (54 °C), for 15 min each at constant shaking. After drying the samples at 60°C overnight, the resulting isolated cell wall residue (CWR) was employed for lignin content determination using the acetyl bromide method (mean extinction coefficient of 23.0772 L g⁻¹ cm⁻¹) (Fukushima and Kerley, 2011) and for lignin compositional analysis using thioacidolysis (Chen *et al*., 2021). The H, G and S lignin monomers released upon thioacidolysis were detected as their trimethylsilyl (TMS) ether derivatives using gas chromatography (GC) coupled to mass spectrometry (MS) based on the specific prominent ions for each compound. The tricin units were quantified after thioacidolysis using high-performance liquid chromatography (HPLC) and the commercial standard of tricin (PHL80987, Sigma). A total of six biological replicates (n=6 per genotype) were analyzed. The differences between genotypes were assessed by t-test with a significance threshold of *P <* 0.05.

### Histochemical tests

WT and *bmr6* internode fragments preserved in 70% ethanol were rehydrated and embedded in 4% agarose. Transverse sections of 100 µm thick were prepared using a vibrating blade microtome (Leica VT1000 S, Leica Biosystems). To detect coniferaldehydes in lignin, sections were incubated in a 2% solution of phloroglucinol prepared in ethanol (w/v) and subsequently acidified with 25% HCl (Blaschek *et al*., 2020). Images were captured after a 5-minute reaction period using a light microscope (Leica LMD7; Leica Microsystems).

### X-ray nanotomography sample preparation

Rehydrated and 4% agarose embedded samples were used to obtain 30 µm sections using a vibrating blade microtome, as previously described. The sections were further dehydrated in a series of increasing concentrations of 30%, 50%, and 70% ethanol for 30 min at each stage. Based on histochemical analysis we extracted the vascular bundles from the second layer below the epidermis, using micro-needles and a stereoscopic microscope (Olympus SZ61). The vascular bundle region was extracted by laser microdissection (Leica LMD7, Leica Microsystems) in a defined area of 30 × 30 µm containing at least three intact tracheary elements. The extracted sample fragments of approximately 30 µm³ were further dehydrated with 80%, 90%, and 100% ethanol to completely remove water before exposure to X-rays. Following, the samples were deposited on a 100 nm thick Si_3_N_4_ membrane (Silson Ltd., UK) for PXCT measurements.

### Ptychographic X-ray computed tomography (PXCT)

The PXCT imaging of tracheary elements were carried out at CATERETE, the coherent scattering beamline at SIRIUS, operated by the Brazilian Synchrotron Light Laboratory (LNLS-CNPEM), Brazil (Meneau *et al*., 2021). For PXCT, the coherent X-ray beam, at 6 keV, passes through a 15 µm pinhole to define the illumination size. The sample positioned 5 mm from the pinhole, was placed on high-precision piezo stages (SmarAct) and was scanned in the *x* and *y* directions with an illumination time of 0.15 s per point. The beam illuminated the sample in a linear trajectory with a step size of 3.15 µm, resulting in an illumination overlap of 79% and satisfying the oversampling requirements for ptychographic image reconstruction (Pfeiffer, 2018). The sample was mounted on an air bearing stage rotational stage (UPR-120 Air-PI), allowing for data acquisition at different angles (Δ_θ_∼150 to 160°). The set of diffraction patterns from each angle produced the angular projections. Since this is an X-ray diffraction-based technique, the patterns (speckles) of each angular projection were recorded in the PiMega 540D detector (LNLS - PiTec, SP, Brazil) placed 28 m from the sample *in vacuum*. The detector has 3072 x 3072 pixels with a 55 x 55 µm^2^ pixel size, resulting in a reconstructed image with a voxel size of 39 nm. The angular scanning range was constrained by the Si_3_N_4_ membrane window’s metallic edges. Three independent regions containing tracheary elements, each derived from a different plant (n = 3 per genotype), were analyzed by X-ray nano-tomography.

### PXCT image reconstruction

Both image projection and tomography of the WT and *bmr6* vascular bundle fragments were reconstructed using advanced algorithms implemented in the pipeline developed by the Scientific Computing Group (GCC) from LNLS (Górecki *et al*., 2023). Diffraction patterns information is retrieved using phase-recovery algorithms, where the diffracted wave is proportional to the Fourier transform of the object (Miao *et al*., 1999). The ptychographic projections were reconstructed using 70 iterations of the relaxed average alternating reflection (RAAR) algorithm and two rounds of 30 and 20 iterations of the alternating projections or the Griffin-Lin (GL) algorithm (Griffin and Lim, 1984; Chang *et al*., 2019). Both algorithms aim to iteratively resolve the phase-retrieval problem until a satisfactory ptychographic image is obtained, in accordance with the experimental data. Finally, 50 iterations of the “*position correction*” algorithm were also added in a second attempt to improve the reconstructions, whose method is based on seeking a correlation between the iterate and the diffraction pattern measured for each scan point (Dwivedi *et al*., 2018). The tomographic reconstructions workflow (Górecki *et al*., 2023) consisted of applying the least squares approach to find shifts and align the frames in the horizontal direction (Prince and Willsky, 1993) and a momentum-based correction to align the vertical direction. The aligned sinogram is used to reconstruct the consistent tomogram using 60 iterations of an expectation-maximization (EM) based method (Fessler, 2000). This iterative process was repeated until convergence was achieved, resulting in a high-resolution three-dimensional reconstruction. The large amount of data required the use of high-performance computers from TEPUI, a computing platform available at LNLS (Furusato *et al*., 2022).

### Image processing and cell wall thickness analysis

Avizo software (Thermo Scientific) was employed to process and analyze the three-dimensional data. First, tomograms were cropped while preserving three complete tracheary elements. The data pre-processing consisted of applying filters to improve segmentation in subsequent stages. Specifically, “nonlocal means” and “unsharp masking” filters were utilized to decrease the noise and highlight the cell wall features of interest within the object. The semi-automatic segmentation was based on pixel intensity threshold, followed by refinement using tools such as “Watershed”, “Magic Wand”, “Remove Islands”, and “Fill Holes”. Following segmentation, the tomograms yielded information on the morphology and thickness of the cell wall. Cell wall thickness measurements were obtained with the “Thickness Map” module, which calculates the diameter of the largest sphere that can be accommodated within the segmented region (Hildebrand and Rüegsegger, 1997). The tracheary elements fragments from independent biological replicates (n=3) per genotype were segmented and the differences between genotypes were assessed by Welch’s t-test with a significance threshold of *P <* 0.05.

### 3D morphometric characterization

The lumen from the previously selected tracheary elements was segmented for morphometric characterization using the Avizo software (Thermo Scientific). For this, a pixel intensity threshold was used as before to pre-select the lumen area, followed by the shrinkage of selected pixels. Once individualized, the area corresponding to an individual cell lumen was independently handled with the “growing’ tool to accommodate all the internal space from that single cell. The created artifacts were excluded using the “lasso” tool. Due to the high resolution, cellular connections (i.e. pit cavities) between adjacent tracheary elements were present and highly visible. The tracheary element lumen was selected in presence and in the absence of these connections. After segmentation, the binary label from the individual cells were exported from Avizo to the FIJI software to determine the tracheary element circularity and convexity parameters (Ménard *et al*., 2022) as follows:

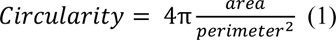

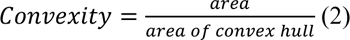

where the convex hull represents the smallest convex polygon that completely encloses the shape. The analysis was applied to all image slices of individual tracheary element lumen length. Each analyzed slice corresponds to a height of 39 nm (the voxel size). In total, three individualized tracheary elements were analyzed per biological replicate (n=9 per genotype). The circularity and convexity values were averaged across all slices for each tracheary element. Each parameter was statistically compared within the genotypes with a two-sample t-test at a confidence *P <* 0.05.

### Numerical simulation of water flow

The surface geometry from the tracheary element lumen, segmented for 3D morphometric characterization for individual cells, was exported from Avizo software as an STL surface file to perform the numerical simulation of water flow. The surface meshes were voxelized onto a regular Cartesian grid for the lattice Boltzmann method (LBM) simulation. The simulations were performed using OpenLB (v1.8-1), an open-source lattice Boltzmann library optimized for parallel computing (Krause *et al*., 2021). The Bhatnagar-Gross-Krook (BGK) collision operator was employed with the D3Q19 lattice model, which uses 19 discrete velocity directions in three dimensions. The fluid (xylem sap) was modelled as water at 20°C with density (ρ) of 998.2 kg m^−3^ and dynamic viscosity (μ) of 1.002 × 10^−3^ Pa s. The characteristic length scale (N) was set in a way that each tracheary element had a simulation voxel size of 0.3 µm. The fluid flow through each segmented tracheary element was driven by a constant axial pressure difference applied between the inlet and outlet boundaries. Simulations were run until a steady state was reached, as confirmed by stabilization of the mean flow velocity. To confirm that the simulations operated within the laminar flow regime, the Reynolds number was calculated for each vessel, remaining well below << 1. Only steady-state velocity and pressure fields were used for the calculation of hydraulic parameters. First, parameters dependent on the driving field were calculated: the lattice Boltzmann method volumetric flow rate (*Q*_*LBM*_) was calculated from the average axial velocity (*v̅Z*) and cross-sectional area (*A*) (Equation 3), and the pressure drop (ΔP) across the vessel segment was determined from the average pressures (*P̅*) at the inlet (*P̅*_*inlet*_) and outlet regions (*P̅*_*outlet*_) (Equation 4).

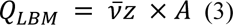

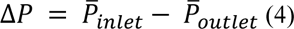

Afterwards, geometric-dependent parameters were computed. Following Ohm’s law analogy for fluid flow (Tyree and Ewers, 1991) the hydraulic resistance (R, Pa s^-1^ m^3^) was calculated (Equation 5).

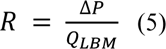

The intrinsic permeability (K, m²) was calculated from Darcy’s law (Equation 6):

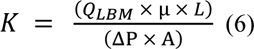

where L is the vessel length.

The simulated hydraulic conductivity (*K*_*n*,*LBM*_, m² Pa^−1^ s^−1^) was calculated based on the Darcy hydraulic conductivity normalized by cross-sectional area (Equation 7).

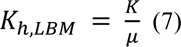

To assess the deviation of real vessel geometry from idealized cylindrical tubes, the *K*_*n*,*LBM*_ was compared with the theoretical hydraulic conductivity calculated by Hagen-Poiseuille (*K*_*n*,*HP*_) (Equation 8):

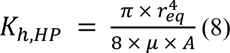

where *r*_*eq*_ is the equivalent radius calculated from the measured cross-sectional area assuming a circular cross-section (Equation 9):

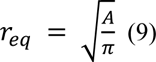

The ratio *Q*_*LBM*_/*Q*_*HP*_ quantifies the hydraulic efficiency of the real vessel geometry relative to an idealized tube with equivalent cross-sectional area. The water flow simulations were applied to three individualized tracheary elements per biological replicate (n=9 per genotype) and each parameter was statistically compared within the genotypes with a two-sample t-test with a significance threshold of *P <* 0.05.

## Results

### The *bmr6* generates shorter sorghum plants

In previous work, we performed a systems biology characterization of the elongating internode of sorghum WT and *bmr6* plants using untargeted metabolomics and large-scale transcriptomics (Ferreira *et al*., 2022). Developing internodes in panicoid grasses show a spatial gradient in cell wall deposition from bottom to top, representing a valuable model system to study secondary wall-related metabolism. To identify a developing internode for these previous experiments, we have cultivated both genotypes under greenhouse conditions and observed that the *bmr6* mutant was slightly shorter than WT by the end of plant development (Ferreira *et al*., 2022), in disagreement with previous studies suggesting that the *bmr6* mutation is normally not associated with detrimental impacts on agronomical traits (Sattler *et al*., 2014). Therefore, here we first evaluated growth parameters from both WT and *bmr6* plants after 140 days of cultivation, when the panicles of all individuals were fully mature (Fig. 1).

**Fig. 1.**
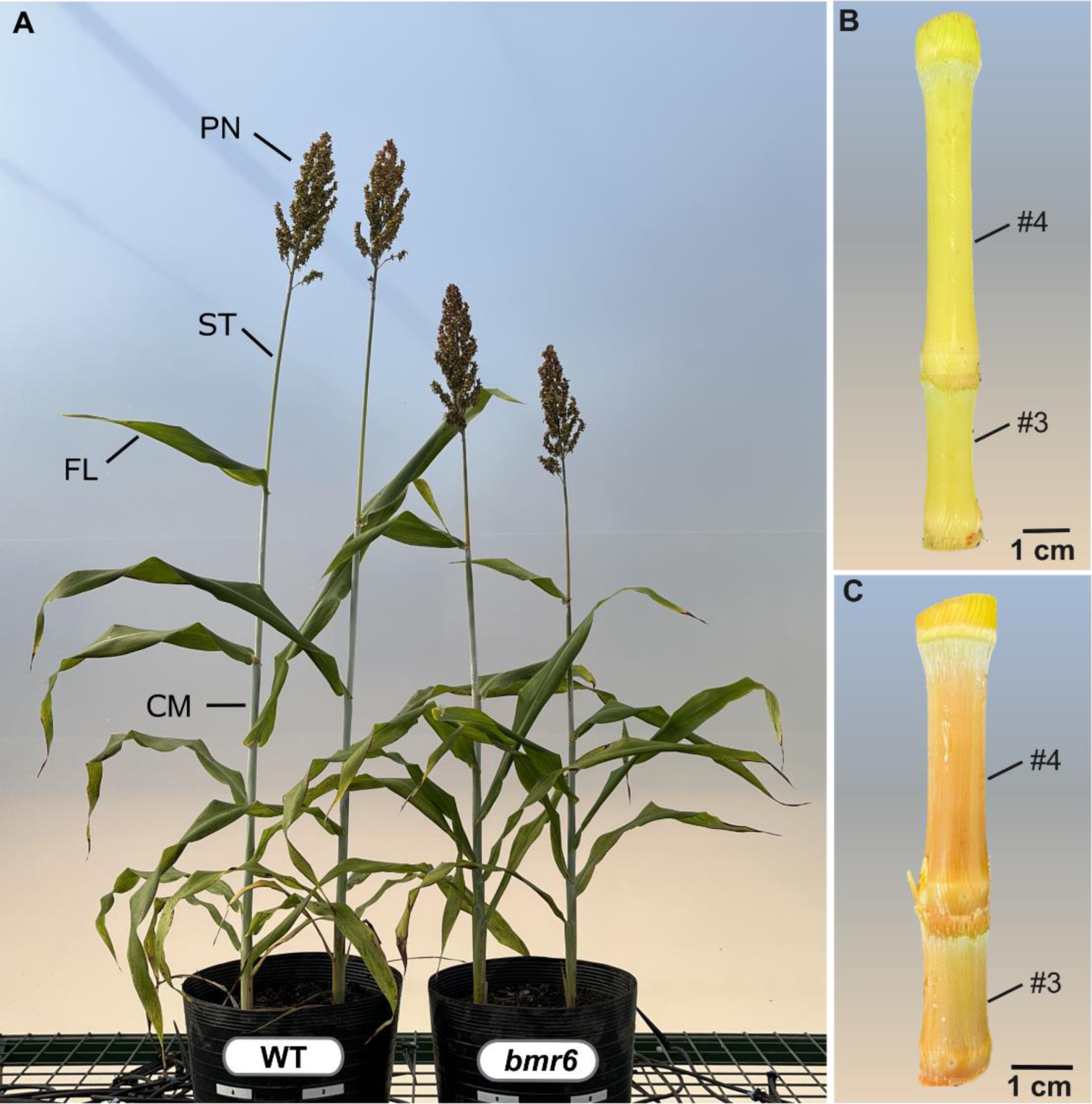
*Sorghum bicolor* material. (A) WT and *bmr6* mature plants. Subsequent experiments were performed using internodes #3 and #4 of WT (B) and *bmr6* (C) plants. Note the brown coloration of *bmr6* mutant internodes. Legend: culm (CM), flag leaf (FL), panicle (PN), and panicle stem (ST).

For *bmr6*, internodes and leaf midveins displayed a typical reddish-brown pigmentation (Scully *et al*., 2016 *b*; Vangala, 2020). Under our growth conditions, the *bmr6* plants were significantly shorter than the WT, whereas no significant differences were found between the genotypes for the other measured parameters, including panicle length, flag leaf length, internode #3 and #4 length and diameter (Table 1). These results are consistent with our previous findings and demonstrate that CAD deficiency did not affect the overall macrostructure of the aerial organs but might result in slightly reduced plant height. Given that (i) plants at this growth stage have finished vegetative development, and (ii) previous solid-state nuclear magnetic resonance (ssNMR) analyses revealed no statistically significant differences in secondary wall deposition between distinct regions of fully mature sorghum internodes (Gao *et al*., 2020), we selected the mid-regions of internodes #3 and #4 for histochemical analysis and PXCT.

**Table 1.** Morphophysiological characterization of WT (n=12) and *bmr6* (n=14) *Sorghum bicolor* plants, cultivated in the greenhouse of the National Laboratory of Biorenewables (LNBr, CNPEM, Campinas, SP).

|  | WT | *bmr6* |
| --- | --- | --- |
| **Plant Height (cm)\*** | 110.59 ± 12.06 | 100.45* ± 8.26 |
| **Panicle Length (cm)** | 18.27 ± 3.63 | 20.54 ± 2.42 |
| **Flag Leaf Length (cm)** | 26.17 ± 6.64 | 30.40 ± 6.77 |
| **Internode #3 Length (cm)** | 2.95 ± 0.99 | 3.78 ± 1.33 |
| **Internode #4 Length (cm)** | 4.64 ± 1.22 | 5.16 ± 1.48 |
| **Internode #3 Diameter (mm)** | 8.49 ± 1.14 | 8.83 ± 1.17 |
| **Internode #4 Diameter (mm)** | 8.05 ± 1.16 | 8.72 ± 1.33 |

\* Significantly different means calculated by $P < 0.05$

### Coniferaldehyde is directly incorporated into lignin

To assess aldehyde incorporation into lignin *in situ*, we applied the Wiesner test (phloroglucinol/HCl) in the internode cross-sections of both WT and *bmr6* plants. In our previous systems biology study, 2D-HSQC (heteronuclear single quantum coherence) NMR on internode samples confirmed that the lignin synthesized by *bmr6* plants is structurally different from that of WT plants, with reduced levels of canonical monolignols and augmented with hydroxycinnamaldehydes (Ferreira *et al*., 2022). However, this bulk analysis performed on whole-cell wall preparations lacks spatial information and fails to provide insights into the distribution of aldehyde incorporation into lignins of distinct cell types. The Wiesner test was shown to react exclusively with coniferaldehyde residues incorporated both at the ends and within lignin polymers (Blaschek *et al*., 2020), allowing specific detection of these residues in different cell types. The sorghum stem exhibits the typical anatomy observed in grasses, characterized by the scattered dispersion of vascular bundles within the fundamental tissue. The rind/Z1 region shows higher density of vascular bundles, which display more lignified sclerenchyma fibers and parenchyma cells compared to the pith/Z2 region (Fig.2).

**Fig. 2.**
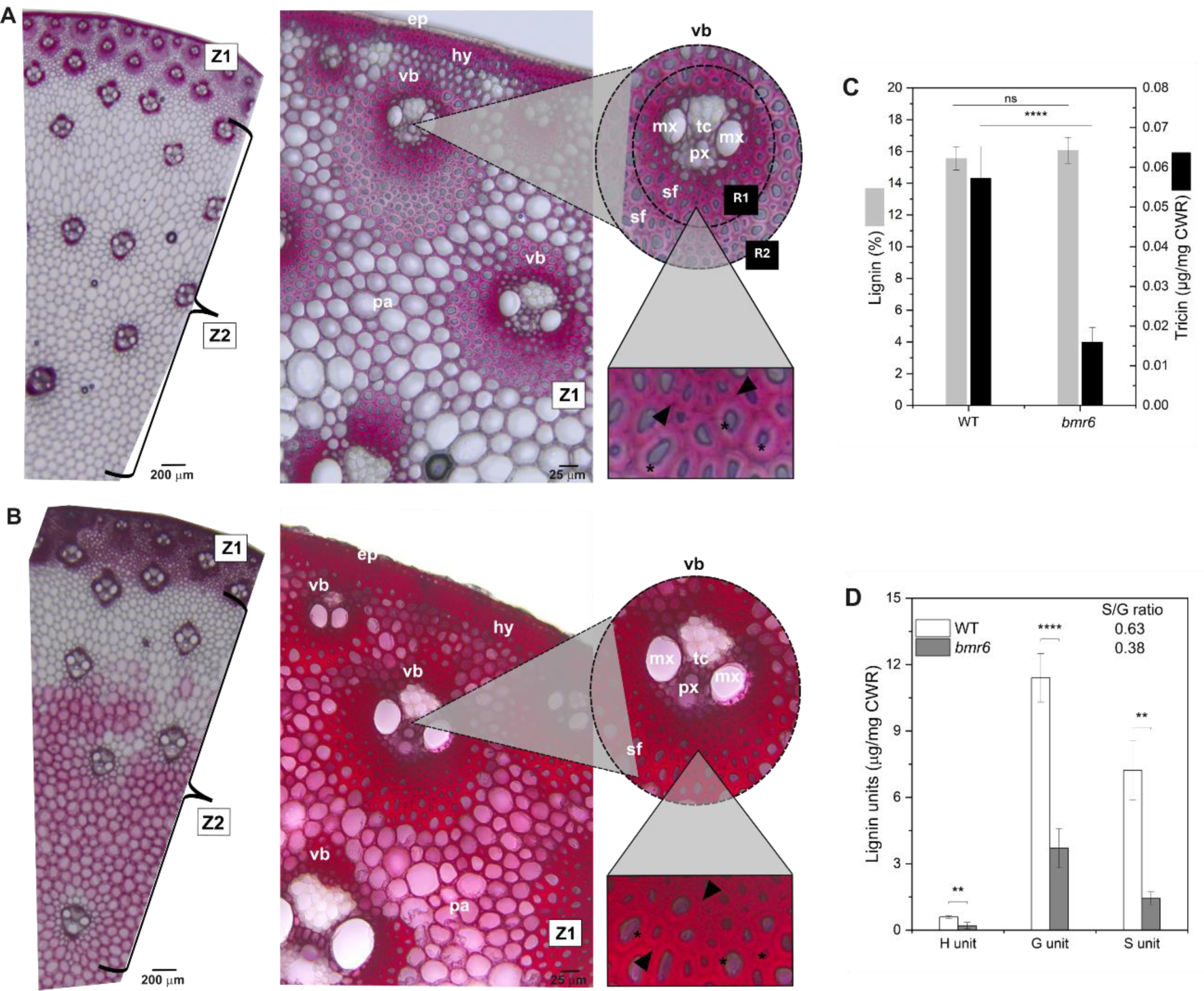
Lignin histochemical and quantitative analyses. *In situ* detection of coniferaldehyde in internode cross sections from WT (A) and *bmr6* (B) stems were performed using the Wiesner test. Positive coniferaldehyde deposition is indicated by the pink/fuchsia coloration. Two main zones are identified: the rind/Zone 1 (Z1) and the pith/Zone 2 (Z2). The central image shows Z1, with small vascular bundles (vb) almost continuously distributed along the culm periphery, in which xylem cells are surrounded by sclerenchyma fibers (sf). Higher magnification of vb (on the top right) shows Regions 1 (R1) and 2 (R2) of the Z1 vascular bundles with identified tissues: epidermis (ep), hypodermis (hy), parenchyma (pa), metaxylem (mx), protoxylem (px) and tracheid (tc). The higher magnification zoom (on the bottom right) from vascular bundles shows the differential deposition of coniferaldehyde residues in the middle lamella and adjacent areas (black arrowhead) and in the secondary cell wall (black asterisks). (C) Total lignin content determined by the acetyl bromide method and quantification of tricin incorporated into lignin determined by thioacidolysis. (D) Lignin composition determined by thioacidolysis. S/G ratio for both genotypes is shown inside the graph. A n=3 per genotype was considered for the statistical analysis and differences are indicated with asterisks: ****, *P <* 0.0001; **, *P <* 0.01; ns, not significant.

Histochemical analysis using phloroglucinol showed that the WT naturally incorporates coniferaldehyde into the lignins of hypodermis (hy), Z1 parenchyma (pa), protoxylem (px), metaxylem (mx), tracheids (tc), and sclerenchyma fibers (sf) (Fig. 2A), indicated by the pink/fuchsia staining, but clear differences in staining intensity were observed among different cell types and even between wall layers of the same cell type. For instance, sclerenchyma fibers located in region 1 (R1), closer to the vascular bundle, exhibited stronger staining compared to fibers in region 2 (R2), which are positioned farther to the xylem and phloem (Fig. 2A, vb higher magnification). We also observed that, in sclerenchyma fibers and hypodermis, the middle lamella and adjacent regions showed stronger staining when compared to secondary cell walls (Fig. 2A, higher magnification zoom). As expected, stronger staining was observed for internode cross-sections of the *bmr6* mutant, indicating increased incorporation of coniferaldehyde residues into lignin (Fig. 2A). Unlike the WT, the *bmr6* also showed staining in the epidermis and in parenchyma cells of the pith (Z2 region). Additionally, *bmr6* plants exhibited homogeneous incorporation of coniferaldehyde in all cell wall layers in both sclerenchyma fibers and hypodermis (Fig. 2B), different from the WT. These results suggest that CAD deficiency results in broad incorporation of aldehydes into the lignin of different cell types and cell wall layers of the same cell type in sorghum.

Given that different results have been reported on whether *bmr6* plants accumulate lower lignin levels than the WT and that the lignin content is one of the major determinants of cell wall architecture and function, we further determined lignin content and composition of internode samples of both WT and *bmr6* prior to PXCT. No significant differences in lignin content were observed between the genotypes (Fig. 2C) whereas the levels of the three canonical monolignols were reduced in the *bmr6* mutant (Fig. 2D). S/G ratio was also strongly reduced in the mutant (Fig. 2D, inset), mainly due to an 80% decrease in the levels of S units. Interestingly, the lignin of *bmr6* also showed significant reduced levels of tricin (Fig. 2C), a flavone characterized as an authentic lignin monomer in grasses. These results confirm that CAD deficiency mainly affects lignin composition and demonstrate that the *bmr6* genotype is a good model to evaluate how changes in lignin structure may affect cell wall architecture and function.

### Higher aldehyde incorporation into lignin has no impact on cell wall thickness

PXCT experiments were carried out to unveil potential morphological and quantitative dissimilarities between the cell walls of WT and *bmr6* internode lignifying cells. The experimental set up is described in Fig. 3A, whereas the detailed parameters for the data collection for each sample are described in Supplementary Table S1. Data collection and analysis focused on a region extracted from the vascular bundle central part, comprising the tracheary elements such as protoxylem and tracheids between the two metaxylem poles (Fig 3B). A total of three tracheary elements regions (n=3), each extracted from different plants per genotype, were sampled for X-ray nano-tomography. The spatial resolution of reconstructed tomographic images ranged from 182 to 246 nm, in the XY plane, and from 168 to 240 nm, in the XZ plane, depending on the data set (Supplementary Table S1), determined by the line profile approach presented in Supplementary Figure S2 and described in the Supplementary Methods. Examples of PXCT reconstructed volumes of approximately ∼30 µm² from tracheary elements from #3 and #4 internodes of WT and *bmr6* tissues are illustrated in Fig. 3B.

**Fig. 3.**
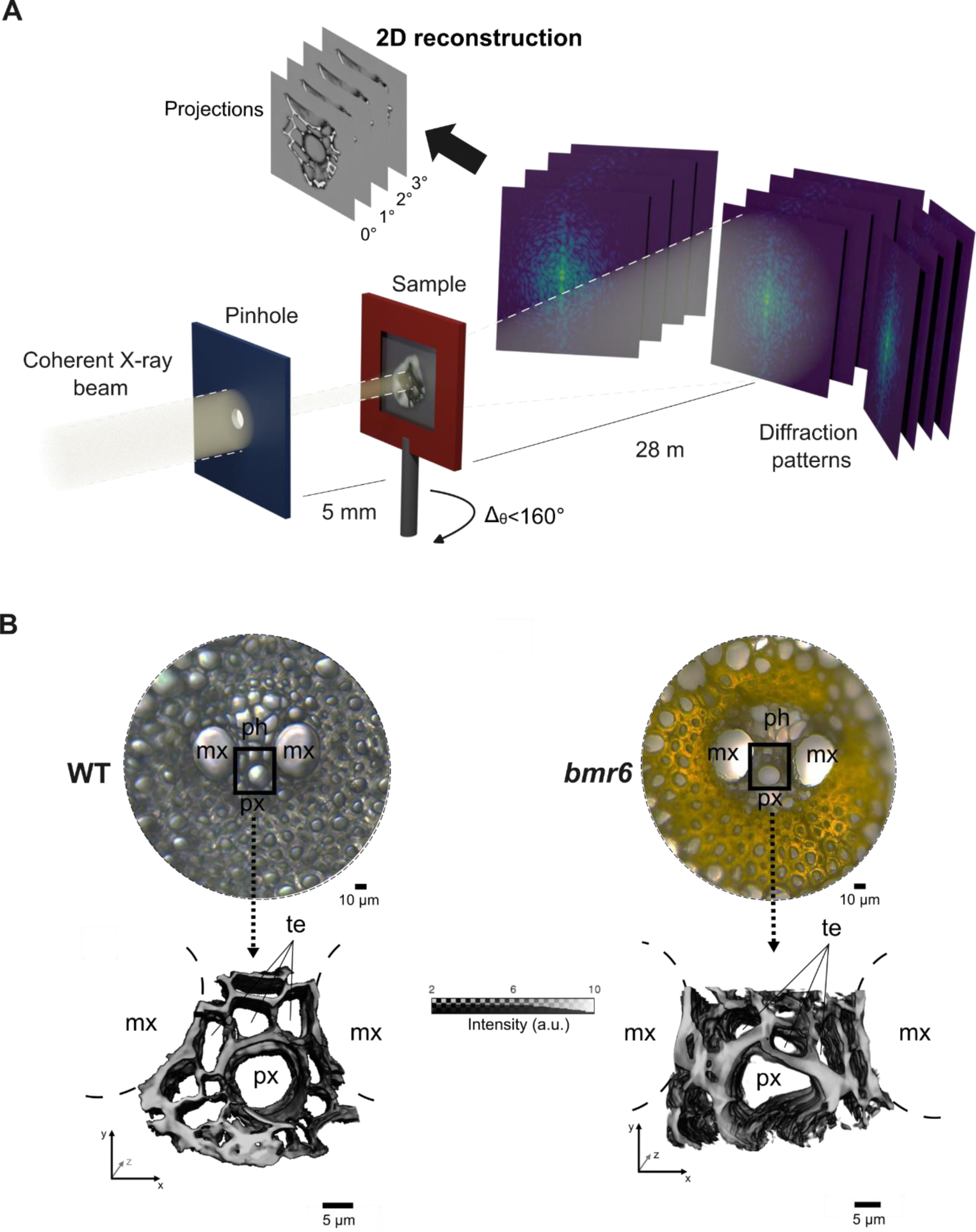
Nano-tomography experiment workflow carried out with sorghum tracheary elements. (A) PXCT experimental setup at CATERETE beamline, at SIRIUS (LNLS-CNPEM). The sample sitting on Si_3_N_4_ membrane is scanned with a coherent beam fraction, whose size is defined by a pinhole. The detector registers the speckle patterns which are used to reconstruct a 2D projection. The three-dimensional data acquisition consists of rotating the sample during the experiment. (B) WT and *bmr6* vascular bundle histology (top). The black box represents the tracheary elements extracted for the PXCT and their respective reconstructed volumes (∼30 µm²) (bottom). Cellular types identified: (mx), protoxylem (px), tracheary elements (te) of the primary xylem and phloem (ph).

Cell wall structure presents heterogeneous morphology depending on the tissue, with varying dimensions along their length, mainly due to naturally occurring intercellular connections. Thus, measuring the wall thickness at a certain height or slice of the cell, as performed in optical and electron microscopies, might not be representative of the real thickness of the cell as a whole. High-resolution PXCT-generated 3D images offer the opportunity for accurate quantitative analysis of cell wall thickness distribution, allowing precise assessment of potential variations along the cell length. The total area was standardized for all the samples based on a normalized length of 12.5 µm and to include three intact cells located at the protoxylem surroundings. The thickness distribution statistics were calculated along the cell height, from each sample, as shown in Fig. 4. Using a stricter wall thickness threshold (>250 nm, image spatial resolution), the averaged median calculated for the WT (1.54 ± 0.17 µm) and *bmr*6 (1.59 ± 0.20 µm) were not statistically different (n=3; Welch’s t-test, P = 0.73). These findings agree with quantitative lignin measurements, demonstrating that CAD deficiency in sorghum had no effects on total lignin content and, consequently, did not affect cell wall thickness distribution along tracheary elements in mature internodes.

**Fig. 4.**
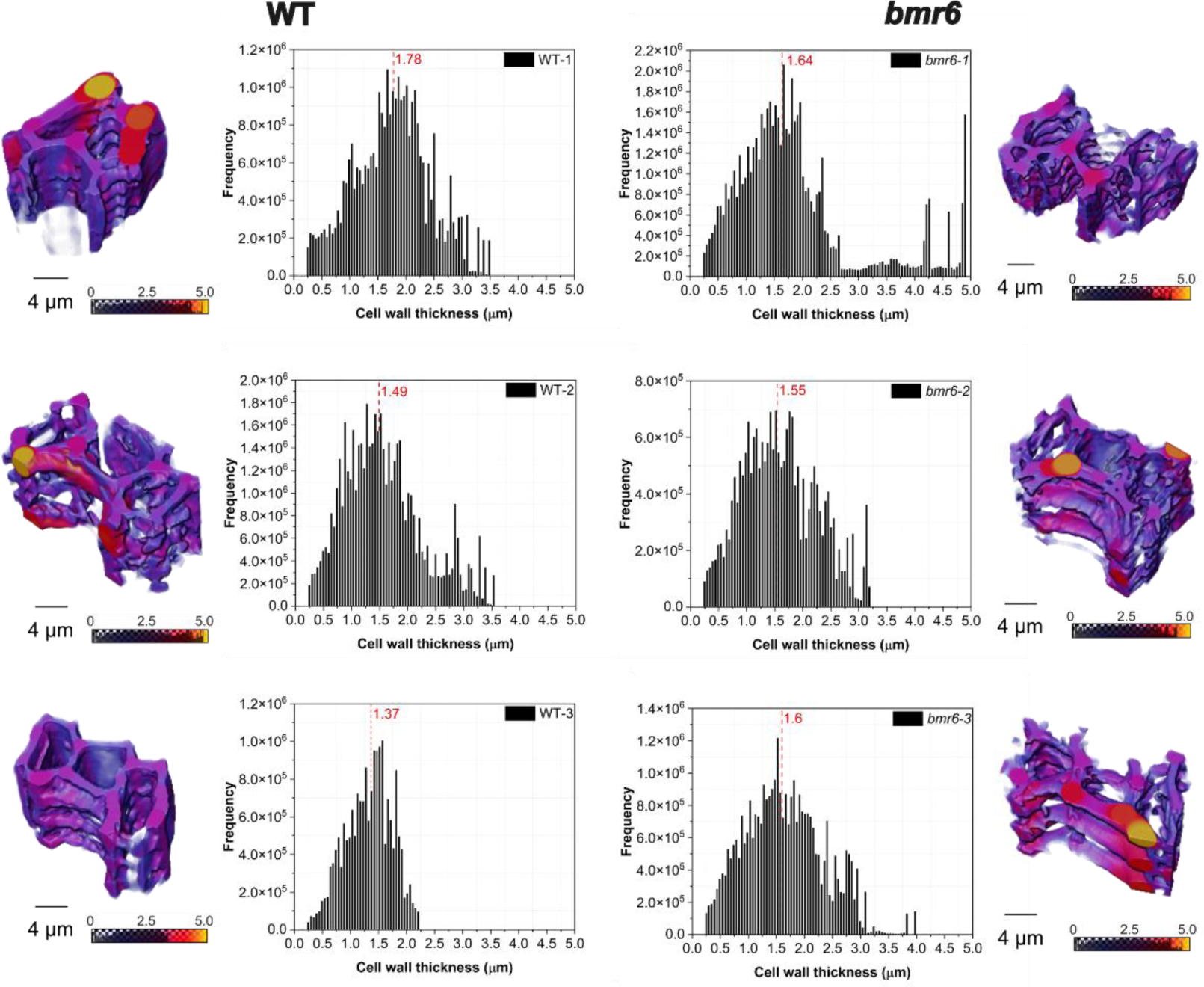
Cell wall thickness distributions map of sorghum internode tracheary elements. The tracheary elements fragments of WT (left side) and *bmr6* (right side) extracted from distinct plants (n=3) per genotype are represented by their color-coded renderings where the darker shades (blue) indicate thinner walls and lighter shades (yellow) represent thicker regions. The corresponding histograms show the thickness distribution highlighting the calculated median values (red dashed line) and statistical differences between genotypes were verified applying a Welch’s t-test with *P <* 0.05.

### Tracheary elements of *bmr6* show reduced convexity, indicative of inward collapse

Precise and spatially controlled lignin deposition is essential for the proper function of xylem water-transporting cells, as lignin confers the required mechanical reinforcement to resist the negative pressure associated with sap transport (Pesquet *et al*., 2025). We benefited from the volumetric information obtained with PXCT to determine circularity and convexity of individual tracheary elements to verify potential morphometric dissimilarities. These parameters depict different aspects of cell morphology: whereas convexity estimates inward collapse, circularity depicts cell deformation (Ménard *et al*. 2022). Measurements of circularity and convexity might help determining the effects of specific lignin chemistries/compositions on mechanical resistance and, consequently, on hydraulics and biomechanics of conducting cells. Previously, such calculations have been performed using 2D measurements of stem cross-sections in different *Arabidopsis* mutants altered in lignin content and/or composition, which allowed mapping single-point cross sectional areas in different tracheary elements (Ménard *et al*. 2022). Here, we took advantage of the 3D data with high spatial resolution to capture potential irregularities along the longitudinal length of single tracheid cells (e.g. lumen twisting, wall thinning along the length, pits, and indentations in the longitudinal axis), which are missing in 2D analyses. The analysis was performed using two different approaches. First, measurements were made in the segmented lumen excluding cellular connections (i.e. pits), thus considering that both cell wall and lumen are continuous structures (Fig. 5A). Accordingly, no significant dissimilarities (*P <* 0.05) were observed between the genotypes, neither for mean calculated circularity (WT = 0.69 ± 0.07; *bmr6* = 0.70 ± 0.05) nor for mean calculated convexity (0.95 for both genotypes) (Fig. 5C). In a second approach, measurements were performed in the segmented lumen with the cellular connections, considering the native non-continuous character from the cellular structures which vary along the longitudinal length (Fig. 5B). When such cellular connections were considered, the mean calculated circularity was not statistically different between the genotypes (0.64 ±0.08 and 0.56 ±0.07, for the WT and *bmr6*, respectively) but the calculated convexity was significantly reduced in cells of the *bmr6* mutant when compared the WT (0.93 ± 0.02 for the WT and 0.89 ± 0.02 for the *bmr6*) (Fig. 5D). These results suggest that the altered lignin composition is associated with changes in collapse resistance of tracheary elements in the *bmr6* mutant when cellular connections are considered.

**Fig. 5.**
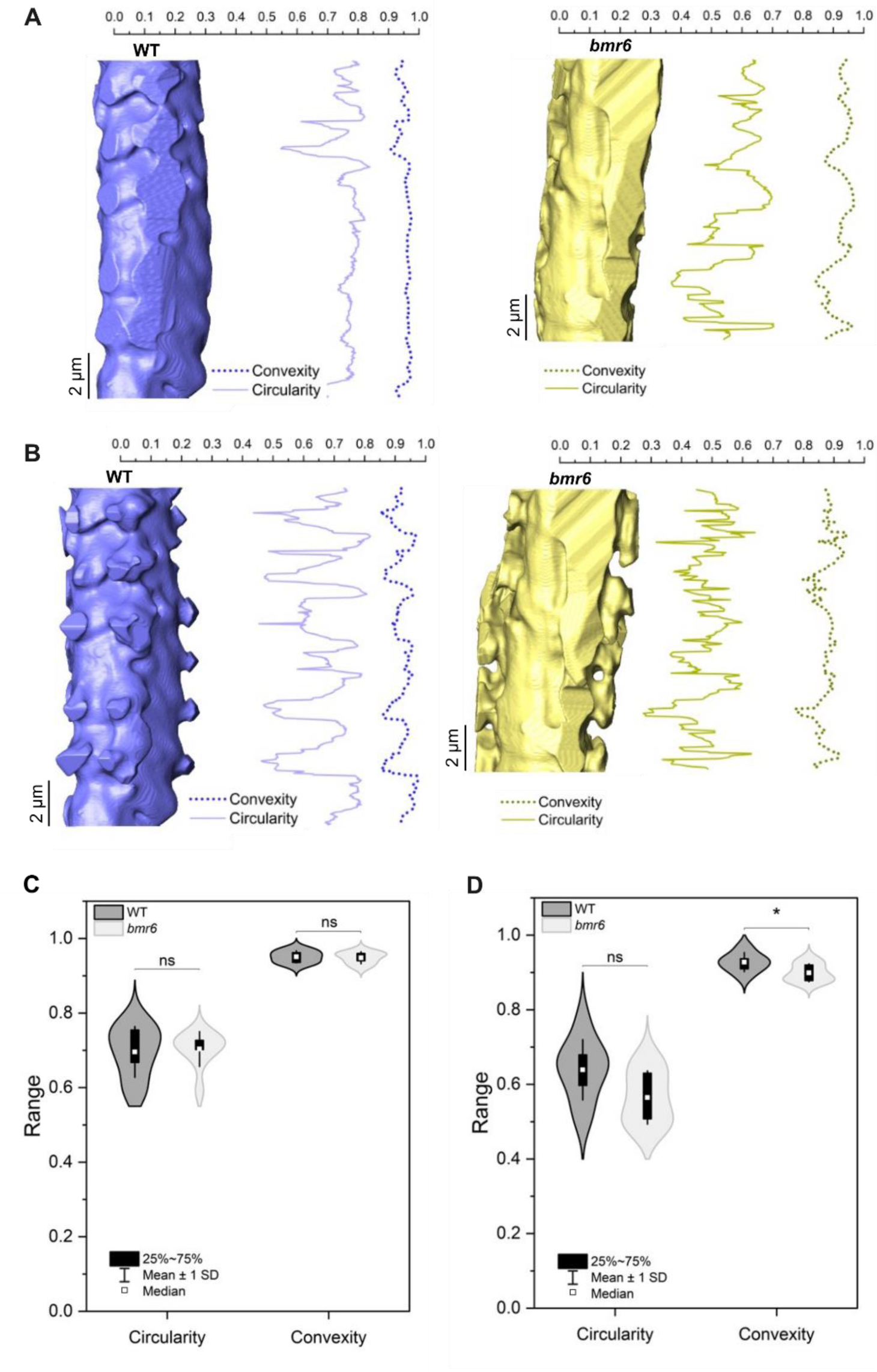
Tracheary elements morphological analysis on three-dimensional images. (A) exemplifies the circularity (continuous line) and convexity (dashed line) analysis done in individual cell lumens of WT (blue) and *bmr6* (yellow) where the cellular connections are disregarded. (B) same analysis, now considering the cell lumens segmented with the cellular connections. C and D, box plots with the circularity and convexity distributions in the absence and in the presence of cellular connections, respectively. A total of three tracheary elements from three biological replicates (n=9) per genotype were statistically compared with a two-sample t-test and no significant (ns) and significant differences (*) are described in the plots considering *P <* 0.05.

### Water flow simulations reveal the contribution of pit-associated lumen geometry

To evaluate whether the subtle geometric differences detected by three-dimensional morphometric analysis result into functional consequences in hydraulic parameters, we performed water flow numerical simulations along independent tracheary elements lumens, which were segmented from PXCT reconstructions. The simulations were conducted using the lattice Boltzmann method under steady-state, laminar flow conditions, with Reynolds numbers well below unity. Velocity field maps revealed the expected parabolic axial velocity profile along cellular lengths for both WT and *bmr6,* with maximum axial velocities concentrated at the lumen center and progressively decreasing toward the cell wall (Supplementary Fig. S3). When the simulations were performed on lumen geometries excluding cellular connections, no significant differences were detected between WT and *bmr6* tracheary elements for permeability, hydraulic resistance, or simulated hydraulic conductivity (Table 2). In both genotypes, the ratio between simulated and theoretical Hagen–Poiseuille conductivity was approximately 0.7, indicating comparable deviation from an idealized cylindrical tube (=1.0). Conversely, inclusion of cellular connections into the lumen geometry depicted genotype-dependent effects. While WT tracheary elements retained similar hydraulic efficiency when the connections are considered, the *bmr6* tracheary elements exhibited a significant reduction in permeability and hydraulic conductivity, with the efficiency ratio decreasing to 0.49 ± 0.12 compared to 0.67 ± 0.10 in WT. These results indicate that, although overall lumen dimensions are preserved in the mutant, the connections associated with geometric features could exert an influence on hydraulic performance.

**Table 2.**
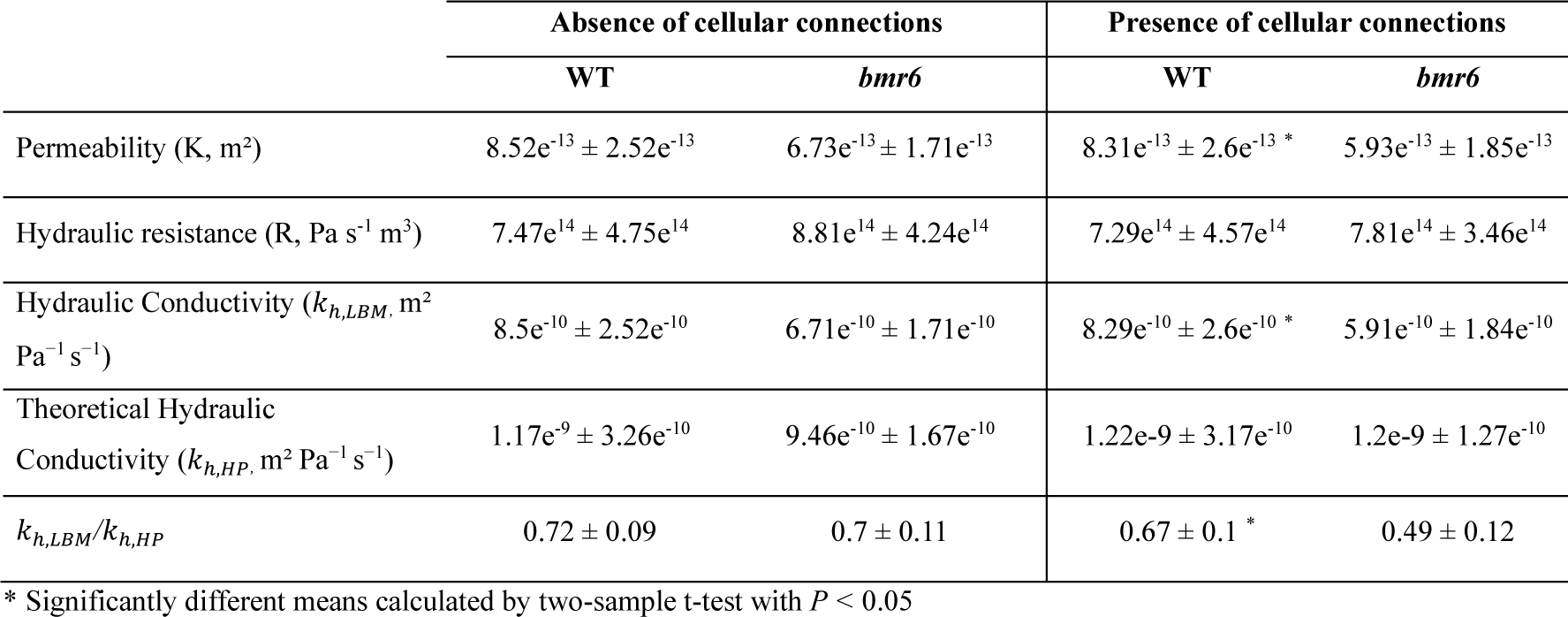
Numerical simulation of water flow through individual tracheary elements reconstructed by PXCT. The parameters were calculated for WT (n=9) and *bmr6* (n=9) lumens with and without cellular connections respectively.

## Discussion

Genetic modifications of lignin biosynthesis have been instrumental in elucidating the structural, mechanical, and physiological roles of lignin in plants, while also offering routes for biotechnological improvement of biomass crops for the bioeconomy. The importance of lignin structures/chemistries in determining cell wall properties and, consequently, in contributing to cell function has been previously undervalued largely due to technical limitations, including lack of genotypes synthesizing lignins with different compositions in phylogenetically diverse species and analytical methods lacking spatial information. In this context, advanced imaging methods with resolution reaching cellular to subcellular levels may allow the proper characterization of the impact of distinct lignin chemistries in cell wall properties and their consequences for cell function. Additionally, new genetic tools allow for the generation of genotypes with altered lignin deposition, which may be targeted to evaluate the contribution of specific lignin structural features. For instance, grass lignins incorporate the flavone tricin as an authentic monomer and contain substantial amounts of *p*-coumarate acylating the side-chains of the polymer backbone (Cesarino *et al*., 2016), but the contribution of these structural features to the biological function of different lignifying cells remains incompletely understood. Here, we took advantage of advanced three-dimensional nano-imaging approaches to investigate how CAD deficiency and the consequent enrichment of hydroxycinnamaldehyde residues into lignin affect xylem cell wall architecture in sorghum.

Consistent with previous studies on the characterization of the *bmr6* mutant (Oliver *et al*., 2005; Palmer *et al*., 2008; Pedersen *et al*., 2008; Sattler *et al*., 2010, 2014; Scully *et al*., 2016; Funnell-Harris *et al*., 2018; Vangala, 2020), the plants displayed a modest but significant reduction in plant height compared with the WT. Yield penalty is a common phenotype observed in plants with altered lignin deposition (Bonawitz and Chapple, 2013), especially when total lignin levels are reduced. However, in our conditions, biochemical assays showed that the lignin content of fully developed internodes of *bmr6* plants was similar to that found in the WT. Apart from lignin reductions affecting the biological function of different cell types, other mechanisms have been linked with plant growth inhibition associated with lignin manipulations. Among them, blocking the monolignol pathway often promotes a rewiring of the phenolic metabolism, which might either prevent the synthesis of growth-promoting compounds or lead to the accumulation of growth-inhibitory/toxic metabolites, resulting in growth inhibition (Muro-Villanueva *et al*. 2019). Indeed, our previous systems biology characterization of the *bmr6* developing internode depicted a massive metabolic reprogramming, and the vast majority of the accumulating compounds were glycosylated (Ferreira *et al*., 2022). As glycosylation is a common mechanism known to inactivate and detoxify harmful compounds (Rates and Cesarino, 2023), these results suggest the activation of detoxification routes for potentially harmful phenolic intermediates that accumulate upon CAD deficiency in sorghum. Given that our PXTC experiments showed reduced tracheary elements convexity in *bmr6*, which might be associated with lower resistance to collapse, we can speculate that the modest reduction in growth observed in this mutant might be a combination of reduced mechanical performance at the whole-plant level and altered soluble phenylpropanoid levels.

Incorporation of aldehyde residues into lignin is known to differ between cell types and cell wall layers, with higher levels found in primary cell walls/middle lamella compared with secondary cell walls of xylem fibers (Hänninen *et al*., 2011; Blaschek *et al*., 2020). Differential incorporation of these units also occurs along wall maturation, as units with terminal aliphatic aldehydes are incorporated at later stages of xylem tracheary elements differentiation (Ménard *et al*. 2022). Our histochemical analyses agree with these observations by demonstrating that coniferaldehyde residues are preferentially found in the middle lamella and adjacent regions when compared to secondary cell walls in sclerenchyma fibers and cells forming the hypodermis in sorghum internodes. Additionally, we noticed a stronger staining in sclerenchyma fibers located closer to xylem conducting cells when compared to fibers more distantly located within the same vascular bundle. The differential incorporation of coniferaldehyde according to cell type and cell wall layer might reflect different mechanical properties demanded by these different cell types/tissues to perform their biological function. Analysis of hypocotyl vasculature in *Arabidopsis* CAD deficient mutants showed that increased aldehyde incorporation into lignin directly influences the hydraulic properties of tracheary elements, causing increased cell wall flexibility and enhancing their capacity to recover from drought (Ménard *et al*. 2022). Molecular dynamic simulations suggest that higher incorporation of coniferaldehyde into lignin changes molecular conformations, torsions and reduces stiffness compared to its corresponding alcohol residues (Ménard *et al*. 2022; Pesquet *et al*. 2025). Coniferaldehyde affects lignin stretching properties by limiting the capacity of the polymer to fold onto itself. Consequently, enhanced coniferaldehyde incorporation might result in lignins with a lamellar extended structure and reduced intra-and intermolecular interactions, directly affecting the hygroscopic capacity and overall biophysical properties of the cell wall (Pesquet *et al*., 2025). As our histochemical analysis showed an increased and more homogenous incorporation of coniferaldehyde along different cell wall layers of *bmr6* lignifying cells, we can speculate that the resulting structural changes found in the lignin polymer contribute to the reduced convexity (and, thus, lower resistance to collapse) observed in tracheids of *bmr6*.

Prior to performing complex and time-consuming PXCT analyses, we have analyzed lignin content and composition to confirm the structural changes caused by CAD deficiency in sorghum internodes. Whereas our results confirm previous data (i.e., unaltered lignin levels, reduced incorporation of canonical monolignols), we observed an additional and hitherto undescribed structural modification: lower levels of tricin incorporated into lignin. Interestingly, the lignin of both rice and maize *cad* mutants contained increased levels of incorporated tricin (Liu *et al*., 2021; Martin *et al*., 2023), although lignocellulosic biomass material evaluated for maize *cad* mutant included a mixture of culms and leaves, which might influence the results as maize leaves naturally incorporates more tricin than culms (Eloy *et al*., 2017). Different tricin levels might also affect lignin structure, as this flavone was proposed to function as nucleation sites that start lignin chains in grasses (Lan *et al*., 2015) and, thus, modification in tricin levels might lead to lignins with altered molecular weight (Li *et al*., 2022). Importantly, such chemical modifications did not translate into overt changes in cell wall thickness at the nanoscale, as we quantitatively assessed cell wall thickness distributions along intact tracheary elements in three dimensions using PXCT and found no statistically significant differences between WT and *bmr6* plants. These results contrast with previous PXCT experiments performed with lignin mutants affected in other lignin biosynthetic genes, such as *CINNAMATE 4-HYDROXYLASE* (*CH4*) in *Arabidopsis*, whose loss-of-function led to altered lignin composition but also reduced lignin content, which ultimately resulted in reduced wall thickness and compromised structural integrity (Polo *et al*., 2020).

The absence of detectable thickness changes despite substantial lignin chemical remodeling highlights the robustness and adaptability of grass secondary cell walls. One plausible explanation is that aldehyde-rich lignin polymers, which have been shown to be less compact and more flexible, may redistribute within the polysaccharide matrix without increasing wall volume. Interactions between lignin, cellulose, and hemicelluloses—particularly xylans that dominate grass cell walls—may buffer changes in polymer chemistry, thereby stabilizing wall dimensions. Compensatory adjustments in cellulose deposition or microfibril organization could further contribute to maintaining constant wall thickness despite altered lignin structure (Hao and Mohnen, 2014; Alemán-Sancheschúlz *et al*., 2020; Shao and Sun, 2024).

Beyond thickness measurements, the three-dimensional nature of PXCT enabled a more nuanced interpretation of xylem elements morphology than is possible with traditional two-dimensional microscopy. By quantifying circularity and convexity along the longitudinal length of individual tracheary elements, we captured subtle geometric features arising from natural irregularities such as pits, longitudinal deformations, and lumen constrictions. While overall circularity values were similar between genotypes, a modest but significant reduction in convexity was detected in *bmr6* tracheary elements when cellular connections were included in the analysis. This observation suggests that CAD deficiency may subtly influence local wall integrity or pit-associated geometry, effects that would likely remain undetected in two-dimensional cross sections. These findings underscore a key advantage of three-dimensional nano-imaging: the ability to evaluate cell wall architecture in its native volumetric context. Metrics such as circularity and convexity, when derived from 3D data, integrate structural variations along the longitudinal axis and provide indirect insight into biomechanical performance. Convexity values close to unity indicate smooth, mechanically stable lumen contours, whereas deviations may signal localized weaknesses or adaptive deformations (Ménard *et al*. 2022; Blaschek *et al*. 2023). Although the observed differences between WT and *bmr6* were subtle and warrant further investigation with larger sample sizes, they demonstrate how volumetric imaging can reveal structure–function relationships that are inaccessible to planar analyses.

From a functional perspective, minor geometric deviations in xylem elements may have implications for hydraulic performance and embolism resistance. The numerical simulation of water flow demonstrated that pit-pair connections in *bmr6* reduce the tracheary elements permeability and simulated hydraulic conductivity. Accordingly, *bmr6* cell exhibited a decrease in the efficiency for hydraulic conductivity when compared to WT. This finding is consistent with the observed decrease in convexity when cellular connections were included in the morphometric analysis and suggests that local deviations from ideal lumen geometry contribute disproportionately to hydraulic resistance. Nonetheless, these results do not imply a loss of vascular function in *bmr6* plants but instead point to a nuanced trade-off between chemical composition and structural organization. The flexible or slightly deformable walls could facilitate accommodation of negative pressures and mitigate drastic collapse under drought stress. In this context, the accumulation of aldehyde-type lignin units may represent an adaptive trade-off: enhancing wall flexibility without compromising overall wall thickness or continuity. Such a mechanism could help explain why *bmr6* plants, despite reduced height, retain functional vascular tissues. The integration of PXCT-derived geometry with numerical flow simulations highlights, once again, the importance of three-dimensional context for interpreting xylem function. Traditional hydraulic resistance and flow are based on idealized cylindrical models (de Araujo *et al*., 2021; Xu and Zhang, 2020; Xu *et al*., 2020, 2023) that disregard lumen irregularities and longitudinal deformations. By directly incorporating experimentally resolved three-dimensional lumen geometries, our simulations demonstrate how small architectural differences—undetectable in two-dimensional analyses—can influence hydraulic behavior. Together, these findings reinforce the view that grass xylem exhibits structural resilience to lignin chemical modification, with functional outcomes also governed by localized geometric features and not only by gross anatomical changes.

In summary, CAD deficiency in sorghum leads to pronounced remodeling of lignin chemistry and spatial deposition patterns without causing major changes in xylem wall thickness. By integrating high-resolution three-dimensional PXCT imaging with biochemical, histological, and computational analyses, this study provides a comprehensive nanoscale perspective on how lignin composition influences cell wall architecture and hydraulic behavior. The results demonstrate that grass xylem exhibits substantial structural resilience to targeted perturbations in lignin biosynthesis, with functional consequences arising primarily from localized geometric features rather than bulk anatomical changes. More broadly, these findings highlight the importance of volumetric nano-imaging approaches for linking cell wall chemistry to structure and function in complex plant tissues.

## Supplementary data

Table S1. Experimental parameters of PXCT data collection and analysis for each sample of WT and *bmr6* (BMR) sorghum plants

Fig. S1. Sorghum bicolor germination process

Fig. S2. PXCT resolution calculation example for the WT-1 plant

Fig. S3. Axial velocity plots examples at different cellular heights for a single cell of tracheary element

Method S1. Image resolution calculation

## Acknowledgements

This research used facilities of the Brazilian Synchrotron Light Laboratory (LNLS), part of the Brazilian Center for Research in Energy and Materials (CNPEM), a private non-profit organization under the supervision of the Brazilian Ministry for Science, Technology, and Innovations (MCTI). We thank LNLS for the provision of PXCT beamtime (proposal number: 20221798) and MSc. Yuri Tonin and Dr. Eduardo X. S. Miqueles, from Scientific Computing Group (GCC) of LNLS, for developing the PXCT code and for their support in image reconstruction.

## Author contributions

LBM, CCP, IC and FFF designed the experiments; LBM, FFF, CCP and LGAL performed the experiments; LBM, FFF, CCP, EM, TK performed the data analysis; CCP, EM, IC, FFF wrote the manuscript draft; CCP, IC and FM reviewed the final manuscript version. All authors read and approved the final manuscript.

## Data availability

The datasets generated and analyzed during the current study are available from the corresponding author on reasonable request.

